# NeoPhAR: Next-generation Phage Therapy for Antimicrobial Resistance

**DOI:** 10.1101/2024.12.27.630320

**Authors:** Sebastián Rivera-Orellana, Justin Yeager, Eduardo Tejera, Andrés López-Cortés

**Affiliations:** Cancer Research Group (CRG), Faculty of Medicine, Universidad de Las Américas, Quito, Ecuador; Biodiversidad Medio Ambiente y Salud, Universidad de Las Américas, Quito, Ecuador; Grupo de Bio-Quimioinformática, Universidad de Las Américas, Quito, Ecuador

## Abstract

Growing antimicrobial resistance (AMR) infections are among the top contemporary concerns in public healthcare systems which can place considerable burdens on healthcare systems. Phage therapy has long been considered a viable option to help combat AMR infections, including on broad geographic scales. One bottleneck in the application of phage therapy is the accurate matching of host-specific phages to bacterial strains, which is traditionally done using molecular techniques. Here we present an open-access deep learning-based model that shows incredible potential to accurately predict phages for therapeutic use. Our goal is to attenuate the matching process so that specific phages, or a cocktail of phages can be prepared for patients with a high probability of therapeutic success, helping to democratize phage therapy treatments even in low to middle-income countries where genomic resources can be limited, costly, or time prohibitive. We feel this is an important first step towards incorporating and applying bioinformatic practices in promising fields such as phage therapy.

## INTRODUCTION

Antimicrobial resistance (AMR) remains one of the most critical challenges to global public health worldwide ^1^. The World Health Organization (WHO) has identified AMR as one of the top 10 global public health threats facing humanity. Infections caused by resistant bacteria, including cefalosporin-resistant *Enterobacteriaceae, Klebsiella pneumoniae*, methicillin-resistant *Staphylococcus aureus* (MRSA), and *Enterococcus faecium* (VRE) impose a significant strain on healthcare systems ^2^. The burden of AMR extends far beyond the number of patients affected. It includes substantial economic costs, with annual expenditures in Europe alone exceeding nine billion euros due to prolonged treatments, extended hospital stays, and high mortality rates. In the United States, the Centers for Disease Control and Prevention (CDC) estimate that AMR infections incur over 20 billion dollars in direct healthcare costs, in addition to 35 billion dollars in lost productivity annually ^3,4^. Acknowledging the gravity of this issue, the 2016 High-Level Meeting of the United Nations General Assembly on Antimicrobial Resistance formally recognized the critical importance of combating AMR. The meeting called on countries to develop and implement their respective National Action Plans, emphasizing the global urgency for coordinated efforts to address this escalating public health crisis ^5^.

AMR infections also contribute significantly to mortality attributed to infectious diseases worldwide. In 2019, it was linked to an estimated 4.95 million deaths globally, surpassing annual fatalities caused by tuberculosis (1.5 million), malaria (643,000), and HIV/AIDS (864,000). Without immediate intervention, global deaths attributable to AMR are projected to reach 10 million annually by 2050. A comprehensive analysis of 204 countries and territories between 1990 and 2021 identified bacterial pathogens as leading causes of sepsis and other severe infectious syndromes. Antibiotic resistance was shown to amplify the risk of death compared to infections caused by susceptible pathogens. These findings underscore the urgent need for a multi-tiered strategy combining preventive and therapeutic approaches to address AMR on a global scale ^1,5^.

Because of this need, there has been renewed interest in integrating phage therapy into the portfolio of treatments targeting AMR ^6^. Phage therapy utilizes bacteriophages—viruses that specifically infect and lyse bacteria—as an alternative or complement to conventional antibiotics. However, a critical step for the efficacy of this approach is accurately identifying host-specific phages that target specific bacterial pathogens. Traditionally, this requires sequencing the pathogen’s genome, an effective but costly and time-consuming process that is often impractical in low-resource settings ^7^.

To democratize and accelerate access to phage therapy, this study introduces a novel deep learning-based approach to predict potential phage-bacteria combinations. The model is trained using a comprehensive database of complete phage genomes and employs neural networks capable of identifying phage-bacteria relationships with high precision. This significantly reduces the time and cost required to determine whether a phage is suitable for treating specific AMR infections. To our knowledge, this is the first approach to combine advanced deep learning and machine learning techniques with complete genomic data to address this challenge in phage therapy.

As a proof of concept, we illustrate how our approach can predict phages relevant to both historical and contemporary infections, providing a robust framework for integrating these predictions into bioinformatics and clinical workflows. Our goal is to empower clinicians to order phage stocks tailored to common infections and update their portfolios periodically in response to emerging needs. This approach represents a significant step toward faster, more precise, and globally accessible phage therapy. Our model demonstrates exceptional predictive performance, achieving high sensitivity and specificity metrics across various bacterial classes. For clinically significant pathogens such as *Mycobacterium, Escherichia, Pseudomonas, Salmonella, Staphylococcus, Klebsiella*, and *Vibrio*, the model exhibited outstanding accuracy. These results highlight its potential to identify effective phages against critical pathogens, making a substantial contribution to the development of alternative therapies for antibiotic-resistant infections.

## METHODS

### Dataset preprocessing

For our prediction model, complete genome data with phage-bacteria information was collected from the PhageAI repository (https://www.phage.ai/), the MillardLab database (https://millardlab.org/), and the NCBI database (https://www.ncbi.nlm.nih.gov/), resulting in a total of 48,500 complete genomes ^8–10^. To curate the database, we selected only phages for which host bacteria information was available, phages with n > 100 for each bacterial genus, and genomes with a minimum nucleotide purity of 95% for ATCG sequences. This filtering process resulted in a total of 37,127 viable phages for this study.

### Molecular descriptors

To obtain molecular descriptors, we used the iLearn tool and server (https://github.com/Superzchen/iLearn?tab=readme-ov-file) for descriptor calculation ^11^. However, repetitive descriptors such as Kmer, DNC, and CKSNAP (1-16), as well as TAC, TCC, TACC, Pse_KNC (65-66), and Pse_DNC (17-18), were identified and excluded from further analysis. Ultimately, we retained 436 unique molecular descriptors, as shown in Table 1.

**Table 1.** Molecular descriptors extracted from bacteriophage genomes.

| Feature Group | Features | Symbol | Number of molecular descriptors calculated |
| --- | --- | --- | --- |
| Nucleic acid composition | Nucleic acid composition | NAC | 4 |
|  | Reverse complement kmer | RCKmer | 10 |
|  | Tri-nucleotide composition | TNC | 64 |
|  | k-spaced nucleic acid pairs | CKSNAP | 96 |
| Electron-ion interaction pseudopotentials | Electron-ion interaction pseudopotentials of trinucleotide | PseEIIP | 64 |
| Autocorrelation and cross-covariance | Dinucleotide-based auto covariance | DAC | 12 |
|  | Trinucleotide-based auto-cross covariance | TACC | 8 |
| Pseudo nucleic acid composition | Pseudo dinucleotide composition | PseDNC | 16 |
|  | Pseudo k-tupler composition | PseKNC | 66 |
|  | Series correlation pseudo dinucleotide composition | SCPseDNC | 28 |
|  | Series correlation pseudo trinucleotide composition | SCPseTNC | 68 |
|  |  | **Total** | 436 |

### Analysis using deep learning

The predictive analysis was performed using a deep learning model specifically designed to identify phages and match them with their host bacteria. This process involved several interconnected stages.

First, the collected data underwent rigorous preprocessing and partitioning. From the 37,127 selected phages and their unique molecular descriptors, the dataset was divided into training (75%) and testing (25%) subsets to ensure robust model evaluation. Prior to partitioning, class balancing techniques were applied, utilizing metrics such as the Euclidean distance between mean vectors to adjust the data and equalize the representation of phages associated with and not associated with each bacterium. The class imbalance was limited to within 5% to maintain data homogeneity.

The deep learning model employed a sequential neural network implemented in Keras 3.0. This network consisted of three dense layers: an initial layer with 16 neurons, an intermediate layer with 8 neurons, and a final output layer with a single neuron. The ReLU activation function was used in the intermediate layers, while a Sigmoid function was applied in the output layer to perform binary classification. Additionally, L1 and L2 regularization techniques, with coefficients of 1e-5 and 1e-4 respectively, were incorporated, along with a Dropout layer with a 50% rate to mitigate overfitting and enhance generalization.

To prevent overfitting and ensure optimal performance, an EarlyStopping mechanism was implemented. This monitored validation loss with a patience threshold of 20 epochs and a minimum improvement criterion of 0.001, stopping the training process if no significant improvements were observed.

An innovative aspect of this analysis was the integration of an application domain space, generated through Principal Component Analysis (PCA). This domain space allowed the identification and exclusion of out-of-range data, improving the model’s accuracy and reliability by ensuring predictions were made only within a relevant domain. PCA also reduced the dimensionality of the molecular descriptors, optimizing computational efficiency and improving model performance. The number of principal components was chosen to capture at least 90% of the explained variance, ensuring an efficient representation of the data. Furthermore, a centroid of the PCA space and a maximum radius, calculated using squared Euclidean distances, were defined to establish the boundaries of the application domain.

Finally, the model’s predictions were applied to new phage data, generating probabilities for the presence of phages associated with each bacterium. These predictions were stored in a CSV file, facilitating subsequent analysis and enabling seamless integration into broader bioinformatics workflows. Loss and accuracy graphs generated during training provided valuable insights into the model’s performance and supported the interpretation of results.

## RESULTS

The findings of this study, which utilized a deep learning approach to predict phages associated with bacteria, demonstrate remarkable predictive performance. The model achieved high accuracy along with consistent sensitivity and specificity metrics, exceeding 85% across various bacterial classes. These results highlight the model’s effectiveness in identifying phages associated with diverse bacteria, representing a significant advancement in biotechnology applied to medicine, particularly in the context of phage therapy (Supplementary Table 1).

### Predictive accuracy

For clinically relevant bacterial pathogens, the model exhibited exceptional performance, achieving outstanding sensitivity and specificity scores for species such as *Mycobacterium* (sensitivity: 0.999, specificity: 0.998), *Escherichia* (sensitivity: 0.899, specificity: 0.892), *Pseudomonas* (sensitivity: 0.934, specificity: 0.923), *Salmonella* (sensitivity: 0.913, specificity: 0.907), *Staphylococcus* (sensitivity: 0.980, specificity: 0.969), *Klebsiella* (sensitivity: 0.945, specificity: 0.944), and *Vibrio* (sensitivity: 0.987, specificity: 0.986). These species are recognized as clinically significant pathogens, and the model effectively predicted phages with potential activity against them—a crucial finding for developing alternative therapies for antibiotic-resistant infections. The high sensitivity and specificity achieved underscore the model’s ability to make accurate classifications while minimizing false positives and false negatives (Supplementary Table 2).

Additionally, the model demonstrated excellent performance with *Streptococcus* (sensitivity: 0.990, specificity: 0.984), *Listeria* (sensitivity: 0.940, specificity: 0.876), and other relevant species, all achieving sensitivity and specificity metrics above 85%. These results emphasize the model’s broad applicability in predicting phages for a wide range of pathogens.

### Predictive ability for low genomic coverage strains

However, for bacteria with limited genomic data availability, such as *Ralstonia* (sensitivity: 0.683, specificity: 0.612) and *Achromobacter* (sensitivity: 0.713, specificity: 0.679), the model exhibited lower performance. This highlights the need for higher-quality and larger datasets to improve prediction accuracy for these species. Nonetheless, these results provide a clear pathway for future improvements, underscoring the importance of expanding available datasets for underrepresented species.

The application of advanced regularization techniques, such as L1 and L2 penalties, combined with the inclusion of a Dropout layer in the neural network, effectively prevented overfitting and ensured the model’s generalizability. Furthermore, dimensionality reduction through PCA enabled the model to efficiently process large volumes of molecular descriptors, enhancing performance without compromising accuracy.

## Discussion

Phage therapy shows incredible promise as an additional tool to help combat AMR infections worldwide. Treatment success however is largely contingent on the accurate matching of phages. Due to clinical and laboratory limitations or considerable delays in the accurate identification of infections, we opted to use a machine learning approach in an attempt to expedite the predictive ability to match phages to likely infection-causing bacterial strains. We have demonstrated high precision in the ability of our model to accurately assign effective phages using whole genome sequence to clinically relevant strains ^12^.

Model predictive accuracy was largely contingent on the completeness of genomic coverage for target bacteria, which is not unexpected. However, we note that even with lower coverage we were able to recover surprisingly high prediction accuracy despite limitations. Given one of our research priorities is to help democratize treatment potentials and develop multi-phage cocktails we feel that the limitations to the model can be easily overcome. Similarly, as genomic databases continue to be updated constantly, the predictive ability of our model should increase in parallel. Our model is also designed to specifically facilitate the ease of blending with other complementary bioinformatic approaches to integrate other open-access methodologies.

Given the heavy contemporary and future estimated, economic burden of AMR infections public health systems would benefit from a diverse portfolio of potential treatments which are widely considered to include phage therapy ^13,14^. Our model can not only be used for patient-specific treatment protocols but also for region-specific AMR monitoring to ensure that sufficient phage production tracks contemporary common infections throughout public and private health systems. Ensuring up-to-date phage production will keep treatment possibilities agile with current demands and reduce costs by streamlining and limiting production to current needs ^15^.

The model developed in this study represents a significant advancement in the integration of bioinformatics and deep learning to address current challenges in antimicrobial therapy. This approach has the potential to revolutionize the treatment of resistant bacterial infections by enabling the personalization of phage cocktails tailored to local epidemiological patterns and the specific genomic profiles of bacteria ^16,17^. Furthermore, its ability to accurately predict the most effective phage combinations could not only optimize the clinical application of these therapeutic agents but also significantly reduce the time required for the design and approval of treatments. Such bioinformatics-driven solutions could play a pivotal role in implementing precision medicine strategies in resource-limited regions, thereby democratizing access to advanced therapies ^18^. Beyond phage therapy, these technologies hold immense promise for designing combination therapies for other infectious diseases, enhancing epidemiological surveillance, and anticipating future bacterial threats. Together, they lay the foundation for a new era in infection management.

As AMR infections continue to increase in frequency, public health systems will greatly benefit from any potential tools that help increase economic treatments with high precision. We have built a model that represents a novel high-precision tool in which to democratize the implementation of phage therapy treatments, including in low and middle-income countries. Not only can the model precisely inform specific cases, but it can be used more broadly to direct phage therapy programs to track current and potential future outbreaks. We feel this is an important contribution to our constantly evolving portfolio of AMR treatment options.

